# Early- but not late-adolescent Western diet consumption programs for long-lasting memory impairments in male but not female rats

**DOI:** 10.1101/2023.10.24.563808

**Authors:** Anna M. R. Hayes, Alicia E. Kao, Arun Ahuja, Keshav S. Subramanian, Molly E. Klug, Jessica J. Rea, Anna C. Nourbash, Linda Tsan, Scott E. Kanoski

## Abstract

Early life Western diet (WD) consumption leads to impaired memory function, particularly for processes mediated by the hippocampus. However, the precise critical developmental window(s) during which WD exposure negatively impacts hippocampal function are unknown. Here, we exposed male and female rats to a WD model involving free access to a variety of high-fat and/or high-sugar food and drink items during either the early-adolescent period (postnatal days [PN] 26-41; WD-EA) or late-adolescent period (PN 41-56; WD-LA). Control (CTL) rats were given healthy standard chow throughout both periods. To evaluate long-lasting memory capacity well beyond the early life WD exposure periods, we performed behavioral assessments after both a short (4 weeks for WD-EA, 2 weeks for WD-LA) and long (12 weeks for WD-EA, 10 weeks for WD-LA) period of healthy diet intervention. Results revealed no differences in body weight or body composition between diet groups, regardless of sex. Following the shorter period of healthy diet intervention, both male and female WD-EA and WD-LA rats showed deficits in hippocampal-dependent memory compared to CTL rats. Following the longer healthy diet intervention period, memory impairments persisted in male WD-EA but not WD-LA rats. In contrast, in female rats the longer healthy diet intervention reversed the initial memory impairments in both WD-EA and WD-LA rats. Collectively, these findings reveal that early-adolescence is a critical period of long-lasting hippocampal vulnerability to dietary insults in male but not female rats, thus highlighting developmental- and sex-specific effects mediating the relationship between the early life nutritional environment and long-term cognitive health.

## 1. Introduction

Consumption of a “Western diet” (WD), characterized as a diet high in saturated fat, refined carbohydrates (including sugars), and “ultra-processed” foods (Cordain et al., 2005), is linked with deleterious effects on both metabolic and cognitive health (Francis & Stevenson, 2013; Heinonen et al., 2014; Shively et al., 2019; Wilson et al., 2007). Increasing evidence indicates that the hippocampus (HPC) is a particularly vulnerable region of the brain to WD-related insults (Davidson et al., 2012; Hsu et al., 2015; Kanoski & Davidson, 2011; Kanoski et al., 2010; Noble et al., 2019). Furthermore, findings from both rodent models and humans reveal that adolescence is a critical period of cognitive development, during which the dietary environment can have long-term implications on cognitive processes (DiGirolamo et al., 2020; Hsu et al., 2015; Larsen & Luna, 2018; Noble et al., 2019; Nyaradi et al., 2014; Pollitt et al., 1995; Tsan et al., 2021; Tsan, Sun, et al., 2022). Although a growing abundance of research indicates a connection between nutrition during adolescence and cognition, adolescence represents a rather long developmental period consisting of ∼9 years in humans and ∼30 days in rats (Sengupta, 2013). Whether there are specific developmental periods within adolescence during which the brain is particularly susceptible to the long-term effects of dietary influences is poorly understood.

During adolescence, the brain undergoes key neurodevelopmental processes linked with pronounced changes in emotion and cognition, including neuronal proliferation, synaptic pruning, myelination, and neuron apoptosis (Georgieff et al., 2015; Konrad et al., 2013; Prado & Dewey, 2014). In addition to being characterized as a period of shifts in social processing and risk behavior, adolescence is also denoted by heightened hippocampal neurogenesis and vulnerability to mnemonic processes (Bayer, 1982; Fuhrmann et al., 2015; He & Crews, 2007; Janssen & Murre, 2008). For example, sugar-enriched high-fat diet (HFD) consumption during the entire adolescent period in rats reduces hippocampal neurogenesis and impairs spatial memory (Boitard et al., 2016). That there may be critical vulnerable windows of development within the adolescent period is suggested by results showing that 1-week HFD consumption from postnatal days (PN) 35-42, which loosely corresponds to mid-adolescence, disrupts doublecortin^+^ (DCX^+^) structure and reduces brain-derived neurotrophic factor (BDNF) expression in the HPC, which are markers of newly proliferated neurons and neural plasticity, respectively (Chiazza et al., 2021). However, these changes were shown to be largely reversed with a 1-week washout on a low-fat diet (Chiazza et al., 2021). It is possible that diet exposure exceeding 1 week in adolescent rats may be required to exert longer-lasting impacts on HPC structure and function. Indeed, the time-course of cell maturation in the HPC reportedly lasts between 1-3 weeks in rodents (Curlik et al., 2014; Kozareva et al., 2019). Given this time-course and that the rate of neurogenesis in the HPC decreases with chronological age (Curlik et al., 2014; He & Crews, 2007), we hypothesize that WD exposure during a ∼2-week window representing early-adolescence will yield long-lasting impacts on HPC-dependent memory processes.

In the present work, we sought to determine whether HPC-dependent memory function is vulnerable to either early-adolescent (15 days from PN 26-41; WD-EA) or late-adolescent (15 days from PN 41-56; WD-LA) exposure to a cafeteria-style WD. Further, to determine whether these distinct adolescent developmental periods differentially program for long-lasting memory impairments, and whether these associations are sexually dimorphic, behavioral assessments were conducted after both short (2-4 weeks) and long (10-12 weeks) periods of healthy diet intervention in both male and female rats. We have previously shown that consumption of this WD model during the entire adolescent period (PN 26-56) leads to HPC-dependent memory impairments that persist despite a 5-week healthy diet intervention in both male and female rats (Hayes et al., 2023; Tsan, Sun, et al., 2022). Here we expand these findings by examining the role of specific developmental windows within the adolescent period (early vs. late) in mediating such effects. To determine whether the dietary influences were independent of obesity status, and specific to HPC-dependent memory processes, we also assessed the potential effects of early vs. late-adolescent WD diet consumption on body weight, body composition, perirhinal cortex-dependent object-novelty recognition memory, anxiety-like behavior, and locomotor activity.

### 2. Materials and Methods

All experiments were approved by the Institutional Animal Care and Use Committee at the University of Southern California (protocol #21096) and performed in accordance with the National Research Council Guide for the Care and Use of Laboratory Animals, which is in compliance with the National Institutes of Health Guide for the Care and Use of Laboratory animals (NIH Publications No. 8023, revised 1978).

#### 2.1. Subjects

A total of 73 male and female Sprague Dawley rats (Envigo, Indianapolis, IN, USA; initial weights of 50-75 g for males and 30-55 g for females) were received at the animal housing facility at the University of Southern California on postnatal day 25 (n=36 males, n=37 females). Animals were singly housed in hanging wire cages with a 12:12 h reverse light/dark cycle in a climate-controlled room (22-24°C, 55% humidity). All animals were maintained on *ad libitum* standard chow (Lab Diet 5001; PMI Nutrition International, Brentwood, MO, USA; 29.8% kcal from protein, 13.4% kcal from fat, 56.7% kcal from carbohydrate) until being assigned to experimental diets (described below) on PN 26.

#### 2.2. Dietary model and experimental design overview

We implemented a “junk food” cafeteria-style diet to model a Western diet (WD) in the present experiments, a model that we previously developed for early life diet exposure in rats (Hayes et al., 2023; Tsan, Sun, et al., 2022) as well as adulthood exposure (Hayes et al., 2022). This WD diet was composed of free-choice *ad libitum* access to high-fat high-sugar chow (HFHS diet; Research Diets D12415, Research Diets, New Brunswick, NJ, USA; 20% kcal from protein, 35% kcal from carbohydrate, 45% kcal from fat), potato chips (Ruffles Original, Frito Lay, Casa Grande, AZ, USA), chocolate-covered peanut butter cups (Reese’s Minis Unwrapped, The Hershey Company, Hershey, PA, USA), and 11% weight/volume (w/v) high-fructose corn syrup-55 (HFCS) beverage (Best Flavors, Orange, CA, USA), placed in separate food/drink receptacles in the home cage for each animal. The 11% w/v concentration of HFCS was selected to match the amount of sugar in sugar-sweetened beverages commonly consumed by humans, as we have previously modeled (Hsu et al., 2015; Noble et al., 2019; Tsan, Sun, et al., 2022). WD rats also had *ad libitum* access to water. Control (CTL) rats received the same number of food/drink receptacles, except that they were filled with only healthy standard chow (LabDiet 5001; PMI Nutrition International, Brentwood, MO, USA; 28.5% kcal from protein, 13.5% kcal from fat, 58.0% kcal from carbohydrate) or water, accordingly.

In order to match initial body weights by group per sex, on PN 26 rats were pseudo-randomly assigned to groups of: early-adolescent consumption of the WD diet (WD-EA, n=12/sex), which was exposure from PN 26-41; late-adolescent consumption of the WD diet (WD-LA, n=12/sex), which was exposure from PN 41-56; or consumption of the healthy standard chow throughout both periods (CTL, n=12-13/sex). These PN periods are based on generalized stages of development in rodents (Sengupta, 2013; Tirelli et al., 2003) and separate the larger adolescent PN 26-56 period we have previously studied with this WD diet model (Hayes et al., 2023; Tsan, Sun, et al., 2022) into distinct early-adolescent and late-adolescent stages. When the WD-EA and WD-LA were not receiving WD, they were given healthy standard chow (LabDiet 5001, the same diet as the CTL group).

Following an ∼2-week re-acclimation period to healthy standard chow after the WD diet exposure for group WD-LA finished on PN 56, which constituted the shorter initial healthy diet intervention period (4 weeks for WD-EA and 2 weeks for WD-LA), a first round of behavioral assessments was performed from PN 71-85. Specifically, Novel Object in Context (NOIC) was performed from PN 71-75, body composition analyses were conducted on PN 78, Novel Object Recognition (NOR) was performed from PN 79-81, Zero Maze was completed on PN 82, and Open Field was conducted on PN 84. A second round of behavioral assessments was performed from PN 125-138, which constituted a longer healthy diet intervention period of 12 weeks for the WD-EA group and 10 weeks for the WD-LA group. For this round of behavior, NOIC was performed from PN 125-129, NOR was conducted from PN 134-135, Zero Maze was completed on PN 136, and Open Field was conducted on PN 138. NOR, Zero Maze, and Open Field were only conducted in females due to our previous findings that 30-d WD diet consumption during the entire adolescent stage in male rats does not affect object-novelty recognition, anxiety-like behavior, or locomotor activity via these assessments (Hayes et al., 2023). A timeline of procedures, which were the same for both sexes, is shown in Fig. 1.

**Figure 1.**
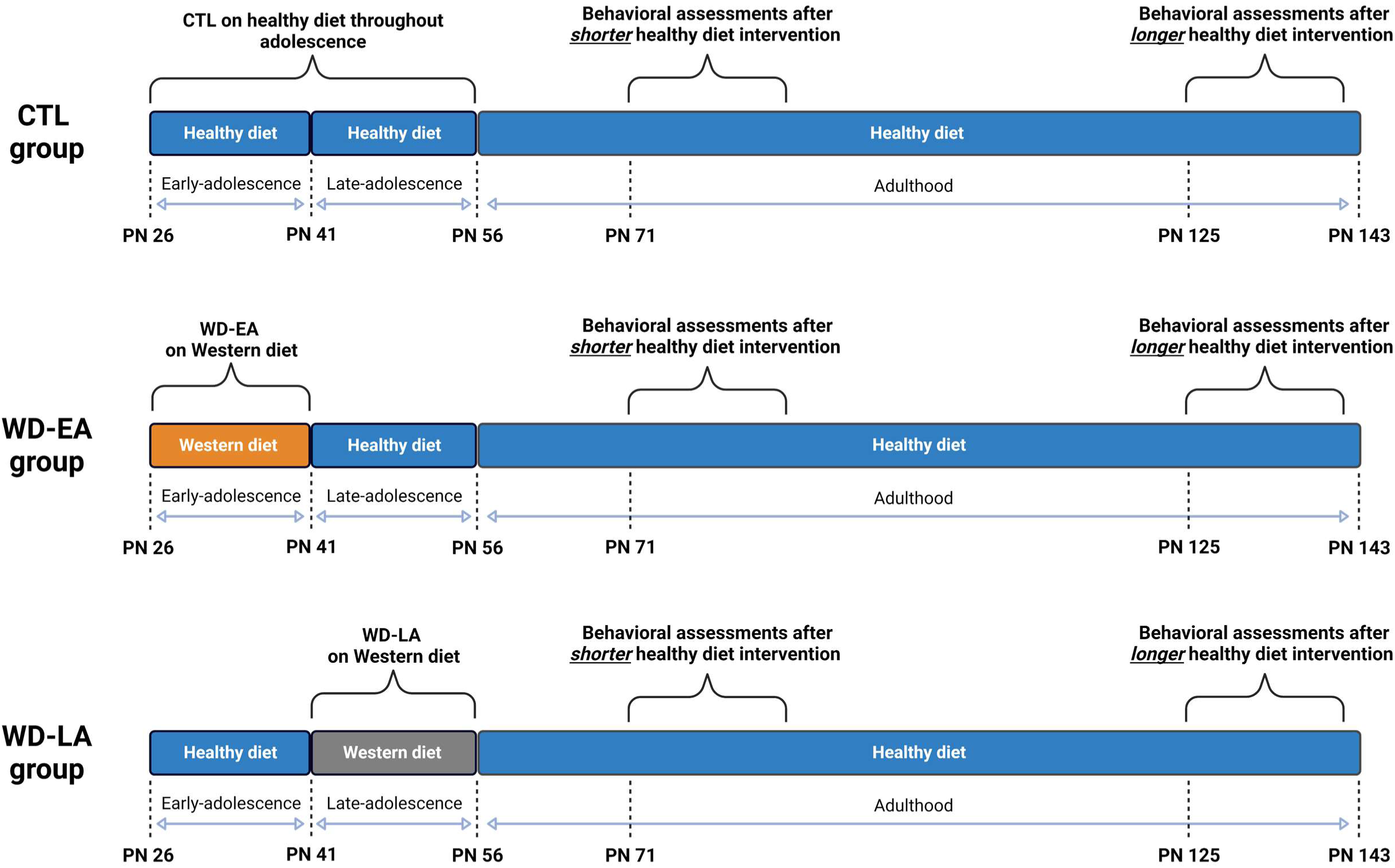
Timeline of experimental procedures for evaluating the effects of early-adolescent (WD-EA) vs. late-adolescent (WD-LA) consumption of a cafeteria-style Western diet (WD), followed by shorter (2-4 weeks) and longer (10-12 weeks) periods of healthy diet intervention in adulthood, on long-lasting memory function and cognition. WD, cafeteria diet; WD-EA, early-adolescent WD diet exposure; WD-LA, late-adolescent WD diet exposure; PN, postnatal day.

#### 2.3. Body weight and energy intake

Body weight and food intake (including spillage collected on cardboard under the hanging wire-bottom cages) were measured three times per week between 8:30-10:30 (shortly before the onset of the dark cycle at 11:00). Total kilocalories consumed from the WD diet components was calculated by multiplying the measured weights of food/drink consumed per rat by the energy density of each component (4.7 kcal/g for HFHSD, 5.7 kcal/g for potato chips, 5.1 kcal/g for peanut butter cups, 0.296 kcal/g for HFCS beverage), and in the same manner the total energy intake for the CTL group was calculated using the energy density of standard chow (3.36 kcal/g). The percentage of total kilocalories consumed from each WD diet components was also calculated using the above values (e.g., kcal from HFHSD / total kcal from all WD diet components). The percentage of kilocalories consumed from each macronutrient (% kcal from carbohydrate, % kcal from protein, % kcal from fat) for the WD diet groups was calculated using the kilocalories consumed from each WD diet component and each component’s macronutrient composition indicated on its nutrition facts label.

#### 2.4. Body composition assessment using nuclear magnetic resonance (NMR)

A Bruker NMR Mini-spec LF 90II (Bruker Daltonics, Inc., Billerica, MA, USA) was used to assess body composition (percent fat mass, percent lean mass, fat-to-lean ratio). Rats were food restricted for 1 h before being weighed and scanned in the Bruker NMR Mini-spec as previously described (Noble et al., 2017), which is based on Time Domain NMR signals from all protons. This system has the benefits of being non-invasive and not requiring anesthesia. Percent fat mass and percent lean mass are calculated as [fat mass (g)/body weight (g)] × 100 and [lean mass (g)/body weight (g)] × 100, respectively.

#### 2.5. Behavioral assessments

##### 2.5.1. Novel Object in Context (NOIC)

NOIC was conducted to assess HPC-dependent contextual episodic memory (Balderas et al., 2008; Martínez et al., 2014). The 5-day NOIC procedure was adapted from previous research (Martínez et al., 2014), and as conducted in previous work from our laboratory (Davis et al., 2020; Hsu et al., 2015; Noble et al., 2021; Suarez et al., 2018; Tsan, Chometton, et al., 2022). Each day consisted of one 5-min session per animal, with cleaning of the apparatus and objects using 10% ethanol between each animal. Days 1 and 2 were habituation to the contexts – rats were placed in Context 1, a semitransparent box (41.9 cm L × 41.9 cm W × 38.1 cm H) with yellow stripes, or Context 2, a black opaque box (38.1 cm L × 63.5 cm W × 35.6 cm H). Each context was presented in a distinct room, both with similar dim ambient lighting yet with distinct extra-box contextual features. Rats were exposed to one context per day in a counterbalanced order per diet group for habituation. Following these two habituation days, on the next day each animal was placed in Context 1 containing single copies of Object A and Object B situated on diagonal equidistant markings with sufficient space for the rat to circle the objects (NOIC day 1). Objects were an assortment of hard plastic containers, tin canisters (with covers), and the Original Magic 8-Ball (two types of objects were used per experimental cohort time point; objects were distinct from what animals were exposed to in NOR). The sides where the objects were situated were counterbalanced per rat by diet group. On the following day (NOIC day 2), rats were placed in Context 2 with duplicate copies of Object A. The next day was the test day (NOIC day 3), during which rats were placed again in Context 2, but with single copies of Object A and Object B; Object B was therefore not a novel object, but its placement in Context 2 was novel to the rat. Each time the rats were situated in the contexts, care was taken so that they were consistently placed with their head facing away from both of the objects. On NOIC days 1 and 3, object exploration, defined as the rat sniffing or touching the object with the nose or forepaws, was quantified by hand-scoring of videos by an experimenter blinded to the animal group assignments. The discrimination index for Object B was calculated for NOIC days 1 and 3 as follows: time spent exploring Object B (the “novel object in context” in Context 2) / [time spent exploring Object A + time spent exploring Object B]. Data were then expressed as a percent shift from baseline as: [day 3 discrimination index – day 1 discrimination index] × 100. Rats with intact HPC function will preferentially explore the “novel object in context” on NOIC day 3, while HPC impairment will disrupt such preferential exploration (Balderas et al., 2008; Martínez et al., 2014).

##### 2.5.2. Novel Object Recognition (NOR)

NOR was used to evaluate exploration of novelty in the form of a novel object, which under testing parameters employed is perirhinal cortex-dependent and not HPC-dependent (Albasser et al., 2011). A grey opaque box (38.1 cm L × 56.5 cm W × 31.8 cm H), placed in a dimly lit room in which two adjacent desk lamps were pointed toward the floor, was used as the NOR apparatus. Procedures followed as described in Noble et al. (2021), modified from Beilharz et al. (2014). Rats were habituated to the empty apparatus and conditions for 10 min 1-2 days prior to testing. The test consisted of a 5-min familiarization phase during which rats were placed in the center of the apparatus (facing a neutral wall to avoid biasing them toward either object) with two identical objects and allowed to freely explore. The objects used were either two identical hard plastic jars with lids or two identical glass vases (first NOR time point), or two identical stemless wine glasses or two identical ceramic vases (second NOR time point). Rats were then removed from the apparatus and placed in their home cage for 5 min. During this period, the apparatus and objects were cleaned with 10% ethanol solution and one of the objects was replaced with a different one (for time point 1: either the jar or vase – whichever the animal had not previously been exposed to – i.e., the “novel object”; for time point 2: either the wine glass or vase – whichever the animal had not previously been exposed to – i.e., the “novel object”). Rats were then placed in the center of the apparatus again and allowed to explore for 3 min. The novel object and side on which the novel object was placed were counterbalanced by treatment group. The time each rat spent exploring the objects was quantified by hand-scoring of video recordings by an experimenter blinded to the animal group assignments. Object exploration was defined as the rat sniffing or touching the object with the nose or forepaws.

##### 2.5.3. Zero Maze

Following established procedures (Noble et al., 2021; Tsan, Chometton, et al., 2022; Tsan, Sun, et al., 2022), the Zero Maze procedure was used to examine anxiety-like behavior. The Zero Maze apparatus consisted of an elevated circular track (11.4 cm wide track, 73.7 cm height from track to the ground, 92.7 cm exterior diameter) that is divided into four equal length segments: two sections with 3-cm high curbs (open), and two sections with 17.5 cm high walls (closed). Ambient lighting was used during testing. Rats were placed in the maze on an open section of the track and allowed to roam for 5 min, during which time they could freely ambulate through the different segments of the track. The apparatus was cleaned with 10% ethanol between rats. The time each rat spent in the open segments of the track (defined as the center of the rat in an open arm) as well as the number of entries into the open segments of the track were measured via video recording using ANYmaze activity tracking software (Stoelting Co., Wood Dale, IL, USA).

##### 2.5.4. Open Field

Open Field was performed to test for locomotor activity as well as anxiety-like behavior following previous procedures (Belzung & Griebel, 2001; Hayes et al., 2022; Noble et al., 2021; Suarez et al., 2018). The Open Field apparatus was a gray arena (53.5 cm L × 54.6 cm W × 36.8 H) with a designated center zone within the arena (19 cm × 17.5 cm). The center zone was maintained under diffused lighting (44 lux) compared to the corners and edges (∼30 lux). For testing, rats were placed in the center of the apparatus and allowed to freely explore for 10 min. The apparatus was cleaned with 10% ethanol between rats. Video recording using ANYmaze activity tracking software (Stoelting Co., Wood Dale, IL, USA) was used to quantify distance travelled in the apparatus during the task (locomotor activity) and time spent in the center zone (a measuring of anxiety-like behavior).

#### 2.6. Statistical Analysis

Data are presented as mean ± standard error of the mean (SEM) for error bars in all figures. Statistical analyses were performed using Prism software (GraphPad, Inc., version 8.4.2, San Diego, CA, USA). Significance was considered at P<0.05. Detailed descriptions of the specific statistical tests per figure panel can be found in Supplementary Table S1. In all cases, model assumptions were checked (normality by the Kolmogorov-Smirnov test and visual inspection of a qq plot for residuals per analysis, equal variance/homoscedasticity by the Brown-Forsythe F test to compare variances and visual inspection of a residual plot per analysis). There were no instances in which normality or homoscedasticity were violated. Group sample sizes were based on previous studies from our lab (Hsu et al., 2015; Noble et al., 2019; Noble et al., 2021) and prior knowledge gained from extensive experience with dietary influences on rodent behavior testing.

### 3. Results

#### 3.1. Neither early-nor late-adolescent WD consumption alters body weight or body composition

Body weight did not differ among male WD-EA, WD-LA, or CTL rats, and all groups exhibited a growth curve (P=0.9147 for main effect of diet, P<0.0001 for main effect of time [indicating growth], and P=0.9990 for interaction of diet × time; Fig. 2A). No differences in body composition were observed for male rats at PN 78 (following WD exposure and after shorter healthy diet intervention period) in terms of percent body fat, fat-to-lean ratio, and percent lean body mass (P=0.3680, P=0.3576, P=0.3351, respectively; Fig. 2B).

**Figure 2.**
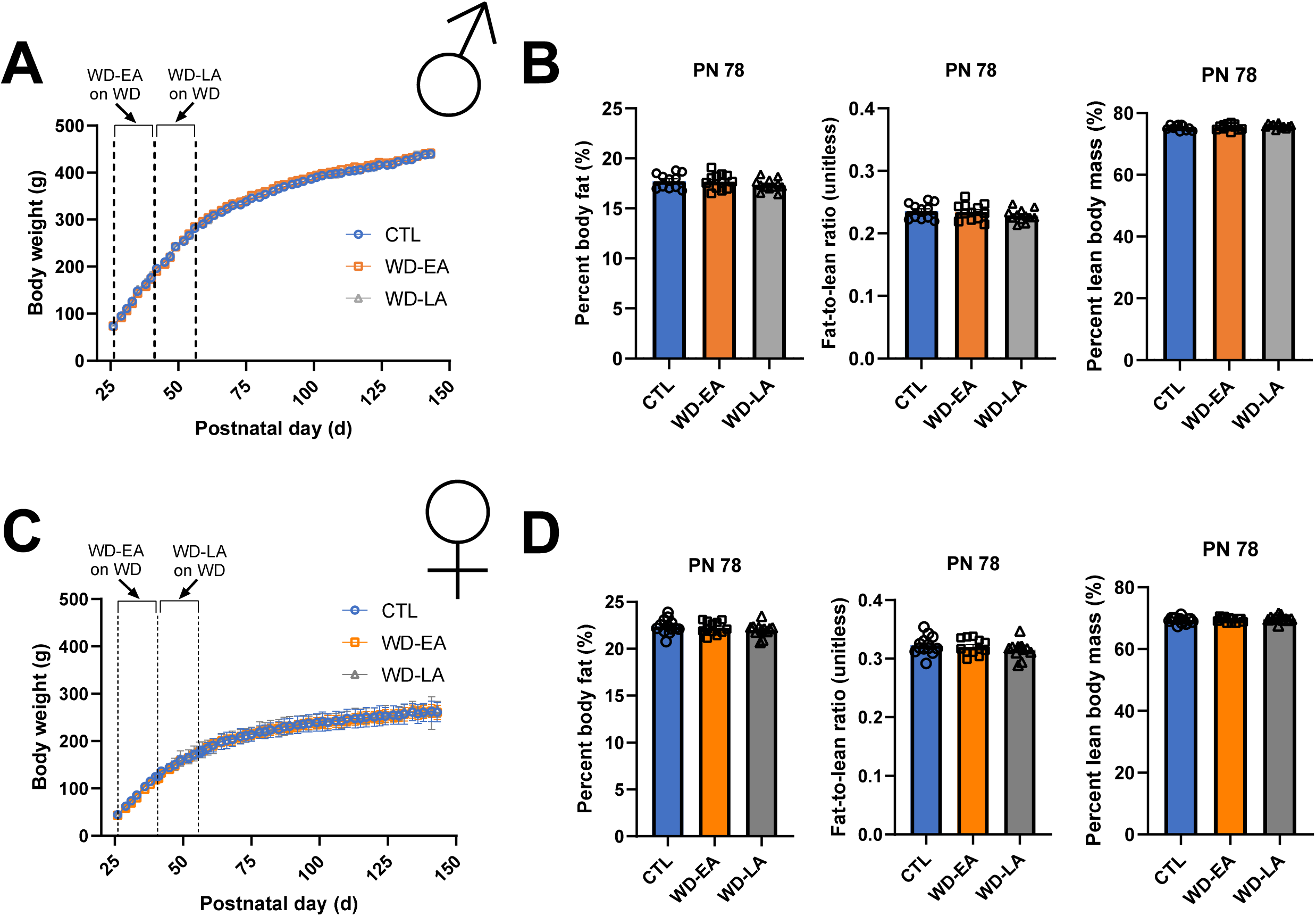
Early-adolescent (WD-EA) vs. late-adolescent (WD-LA) consumption of a cafeteria-style Western diet (WD) does not significantly alter body weight trajectories or body composition in male or female rats. (A) Body weight over time for male rats, with periods of exposure to the WD diet indicated within the dotted vertical lines (two-way ANOVA with diet, time [as repeated measure], and a diet × time interaction as factors; for diet, P=0.9147; for time, P<0.0001; for diet × time interaction, P=0.9990). (B) Body composition for male rats at PN 78 after the shorter healthy diet intervention period (one-way ANOVA with diet as the factor; % fat, P=0.3680; fat-to-lean ratio, P=0.3576; % lean, P=0.3351). (C) Body weight over time for female rats, with periods of exposure to the WD diet indicated within the dotted vertical lines (Mixed effects model with factors of diet [fixed], time [fixed, as repeated measure], diet × time interaction [fixed], subject [random]; for diet, P=0.9999; for time, P<0.0001; for diet × time interaction, P=0.8673). (D) Body composition for female rats at PN 78 after the shorter healthy diet intervention period (one-way ANOVA with diet as the factor; % fat, P=0.4590; fat-to-lean ratio, P=0.8414; % lean, P=0.3947). Error bars represent ± standard error of the mean; n=11-13/group. ANOVA, analysis of variance; WD, cafeteria diet; WD-EA, early-adolescent WD diet exposure; WD-LA, late-adolescent WD diet exposure; PN, postnatal day.

For female rats, body weight did not differ among WD-EA, WD-LA, or CTL rats, and all groups exhibited a growth curve (P=0.8673 for main effect of diet, P<0.0001 for main effect of time [indicating growth], and P=0.9999 for interaction of diet × time; Fig. 2C). Female rats did not differ in body composition at PN 78 in terms of percent body fat, fat-to-lean ratio, and percent lean body mass (P=0.4590, P=0.8414, P=0.3947, respectively; Fig. 2D).

#### 3.2. WD consumption during either early-or late-adolescence increases energy intake

Total energy intake (kcal/24 h) in male rats significantly differed at certain time points between PN 31-61 among the WD-EA, WD-LA, and CTL groups, although there was no significant main effect of diet (P=0.6394 for main effect of diet, P<0.0001 for main effect of time, and P<0.0001 for interaction of diet × time; Fig. 3A). In general, the group being given the WD diet during its respective WD window occasionally ate significantly more kcal than the other two groups being given standard chow (e.g., WD-EA rats consumed more kcal on PN 40 while on WD diet than WD-LA and CTL rats, and WD-LA rats consumed more kcal than WD-EA and CTL rats on PN 45; Fig. 3A; see Supplementary Table S1 for all post hoc comparisons). There were no statistically significant differences between percentage of kcal consumed from the various WD components between the male WD-EA and WD-LA groups (Fig. 3B; Supplementary Fig. S1A). Furthermore, there were no differences in percentage of kcal consumed from fat, carbohydrate, or protein between male WD-EA vs. WD-LA rats (Fig. 3B).

**Figure 3.**
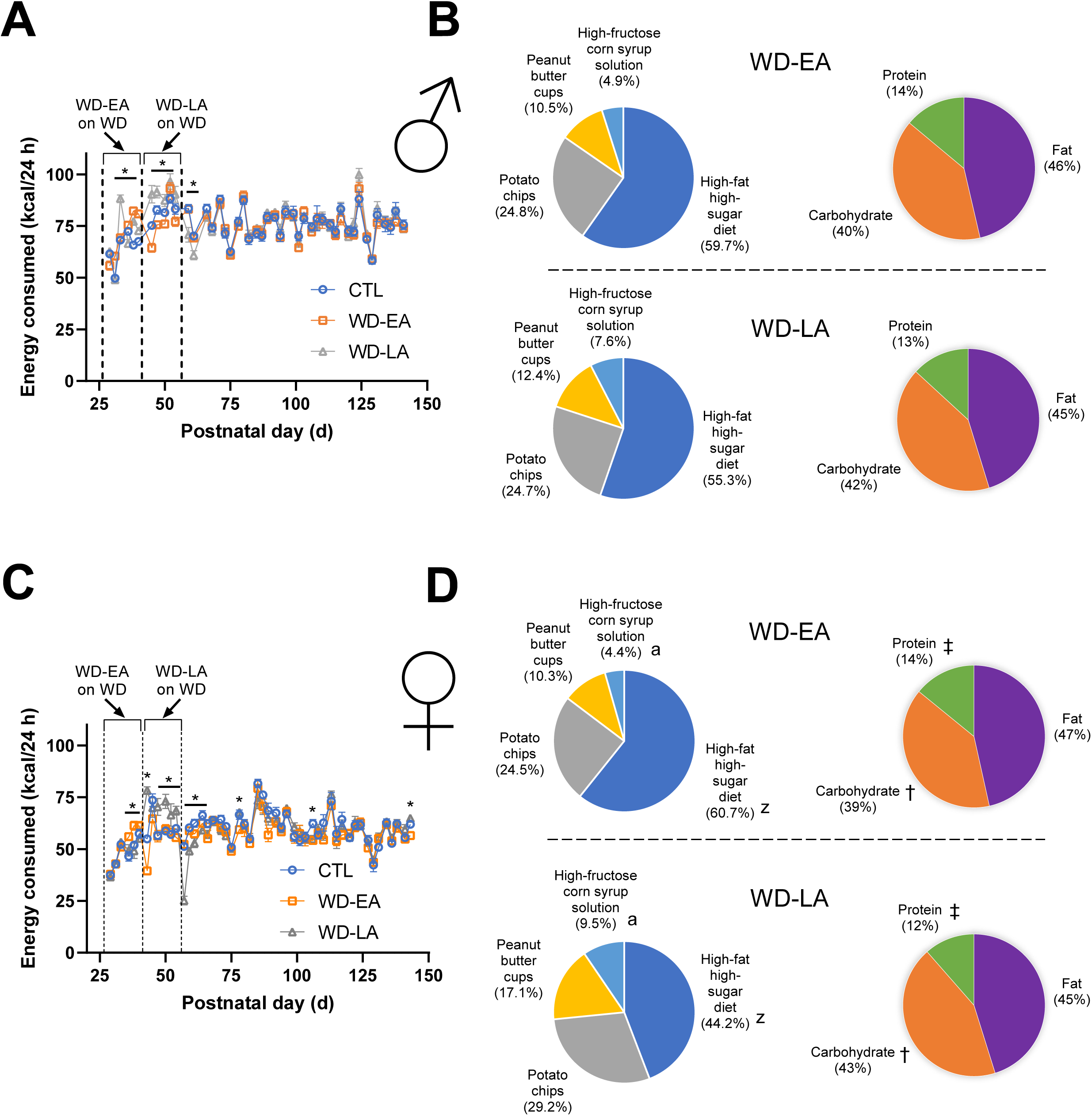
Early-adolescent (WD-EA) vs. late-adolescent (WD-LA) consumption of a cafeteria-style Western diet (WD) increases energy intake during the WD diet period in male and female rats. (A) Energy consumption over time in male rats, with periods of exposure to the WD diet indicated within the dotted vertical lines (two-way ANOVA with diet, time [as repeated measure], and a diet × time interaction as factors; for diet, P=0.6394; for time, P<0.0001; for diet × time interaction, P<0.0001 – post hoc differences listed in Supplementary Table S1). (B) Percentage of kcal consumed from the WD diet components and percentage of kcal consumed per macronutrient for male WD-EA and WD-LA groups (unpaired t-test for all except the % kcal from peanut butter cups comparison, for which a Mann Whitney test was used; % kcal from HFHSD: P=0.5030, % kcal from potato chips: P=0.8058, % kcal from peanut butter cups: P=0.9323, % kcal from HFCS beverage: P=0.1358, % kcal from fat: P=0.2983, % kcal from carbohydrate: P=0.0995, % kcal from protein: P=0.3855). (C) Energy consumption over time in female rats, with periods of exposure to the WD diet indicated within the dotted vertical lines (Mixed effects model with factors of diet [fixed], time [fixed, as repeated measure], diet × time interaction [fixed], subject [random]; for diet, P=0.3247; for time, P<0.0001; for diet × time interaction, P<0.0001 – post hoc differences listed in Supplementary Table S1). (D) Percentage of kcal consumed from the WD diet components and percentage of kcal consumed per macronutrient for female WD-EA and WD-LA groups (unpaired t-test for all except the % kcal from carbohydrate comparison, for which Welch’s test for unequal variances was used; % kcal from HFHSD: P=0.0068, % kcal from potato chips: P=0.2869, % kcal from peanut butter cups: P=0.1188, % kcal from HFCS beverage: P=0.0055, % kcal from fat: P=0.2212, % kcal from carbohydrate: P=0.0018, % kcal from protein: P=0.0037). Error bars represent ± standard error of the mean; n=11-13/group; *P<0.05, ^a^ P<0.01 % kcal from HFCS comparison between female WD-EA and female WD-LA, ^z^ P<0.01 % kcal from HFHSD comparison between female WD-EA and female WD-LA † P<0.01 % kcal from carbohydrate comparison between female WD-EA and female WD-LA, ‡ P<0.01 % kcal from protein comparison between female WD-EA and female WD-LA. ANOVA, analysis of variance; HFCS, high-fructose corn syrup; HFHSD, high-fat high-sugar diet; WD, cafeteria diet; WD-EA, early-adolescent WD diet exposure; WD-LA, late-adolescent WD diet exposure; kcal, kilocalories; PN, postnatal day.

In females, similar to males, total energy intake (kcal/24 h) significantly differed at certain time points between PN 36-66 among the WD-EA, WD-LA, and CTL groups, but there was no significant main effect of diet (P=0.3247 for main effect of diet, P<0.0001 for main effect of time, and P<0.0001 for interaction of diet × time; Fig. 2C). When a group was given WD diet, they consumed more kcal in general than the other two groups being given standard chow at the time (e.g., WD-EA rats consumed more kcal on PN 38 while on WD diet than WD-LA and CTL rats, and WD-LA rats consumed more kcal than WD-EA and CTL rats during PN 47-54; Fig. 3C; see Supplementary Table S1 for all post hoc comparisons). The increase in energy consumed was generally more pronounced for WD-LA (∼20% increase from CTL) than WD-EA (∼16% increase from CTL). In terms of percent kcal breakdown for females, WD-EA rats consumed a significantly smaller percentage of kcal from the high-fructose corn syrup solution than WD-LA rats (P=0.0055), whereas WD-LA rats consumed greater kcal from the high-fat high-sugar diet than WD-EA rats (P=0.0068; Fig. 3D; Supplementary Fig. S1B). WD-EA rats consumed a smaller percentage of kcal from carbohydrate (39%) compared to WD-LA rats (43%, P=0.0018), and as a result 2% higher percentages of kcal were consumed from protein and fat for WD-EA relative to WD-LA, although this difference was only statistically significant for the protein comparison (P=0.0037 for % kcal from protein, P=0.2212 for % kcal from fat; Fig. 3D).

#### 3.3. Both early- and late-adolescent WD consumption impairs HPC-dependent memory function in male and female rats after shorter-term healthy diet intervention

Male and female WD-EA and WD-LA rats exhibited impairments in NOIC after the shorter healthy diet intervention period (P=0.0002 overall for males, P=0.0017 overall for females), manifested as percent shifts from baseline object exploration that were significantly different from CTL rats and did not differ from zero (Fig. 4; see figure caption and Supplementary Table S1 for post hoc statistical comparisons). These NOIC results were consistent when examined as discrimination indices on day 3 and day 5 of the NOIC task (Supplementary Fig. S2; i.e., no differences in baseline discrimination index for the future novel object but differences in discrimination index on the test day for the time point after the initial shorter healthy diet intervention period), and occurred in absence of any differences in total object exploration time on day 3 and day 5 of the NOIC task (Supplementary Fig. S2).

**Figure 4.**
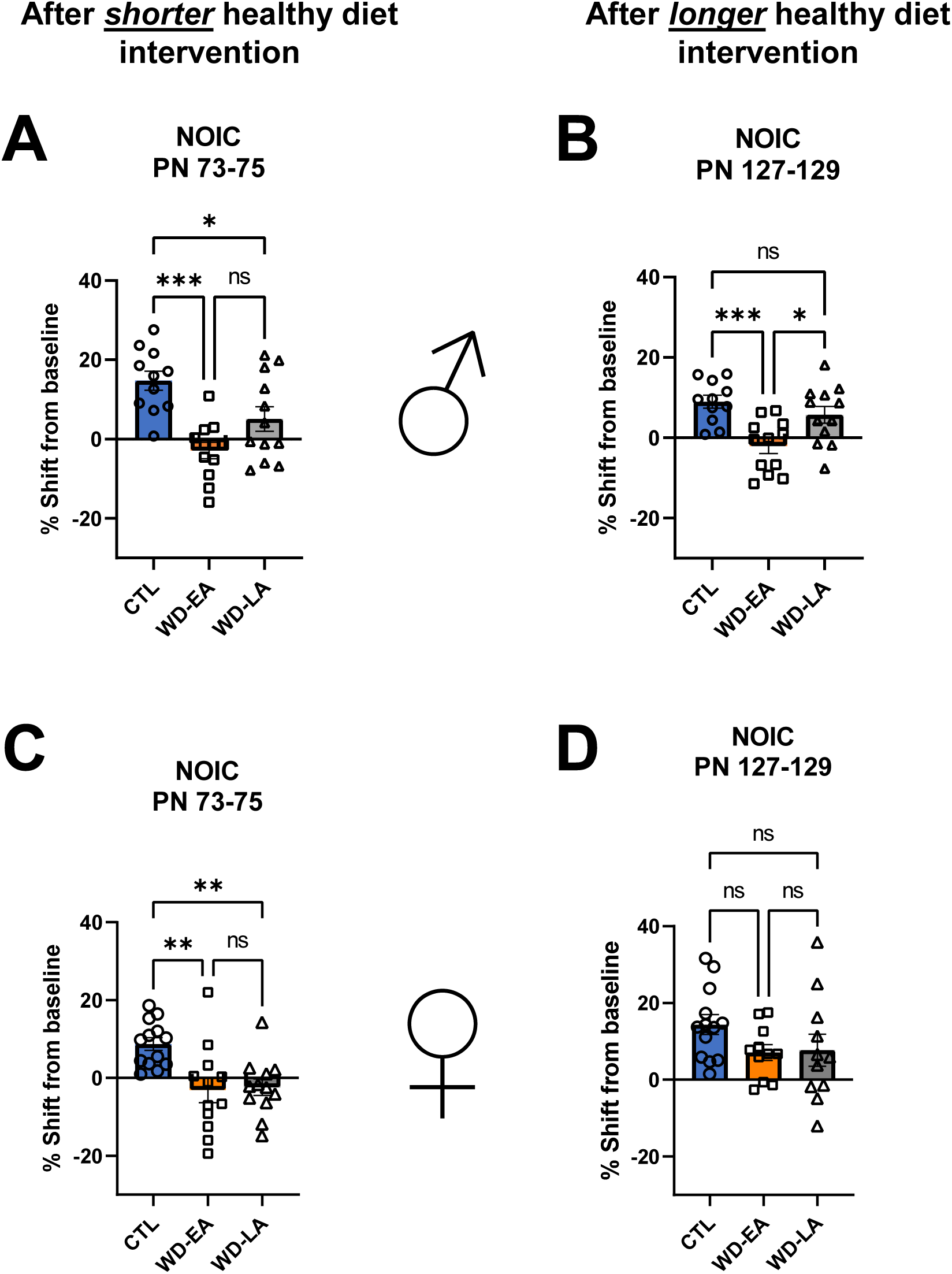
Sex-specific effects of early- or late-adolescent WD diet consumption (WD-EA) on long-lasting hippocampal-dependent memory function. (A) Hippocampal-dependent memory function assessed via NOIC and expressed as percent shift from baseline object exploration in male rats after early-adolescent (WD-EA) and late-adolescent (WD-LA) WD diet exposure and after the initial shorter healthy diet intervention period (one-way ANOVA with diet as the factor; P=0.0002 overall; post hoc Tukey’s multiple comparisons test: CTL vs. WD-EA, P=0.0001; CTL vs. WD-LA, P=0.0331; WD-EA vs. WD-LA, P=0.0840). (B) Hippocampal-dependent memory function in male rats after the longer healthy diet intervention period (one-way ANOVA with diet as the factor; P=0.0002 overall; post hoc Tukey’s multiple comparisons test: CTL vs. WD-EA, P=0.0007; CTL vs. WD-LA, P=0.4541; WD-EA vs. WD-LA, P=0.0146). (C) Hippocampal-dependent memory function in female rats after early-adolescent (WD-EA) and late-adolescent (WD-LA) WD diet exposure and after the initial shorter healthy diet intervention period (one-way ANOVA with diet as the factor; P=0.0017 overall; post hoc Tukey’s multiple comparisons test: CTL vs. WD-EA, P=0.0041; CTL vs. WD-LA, P=0.0073; WD-EA vs. WD-LA, P=0.995). (D) Hippocampal-dependent memory function in female rats after the longer healthy diet intervention period (one-way ANOVA with diet as the factor; P=0.1732 overall). Error bars represent ± standard error of the mean; n=11-13/group; *P<0.05, **P<0.01, ***P<0.001. ANOVA, analysis of variance; WD, cafeteria diet; WD-EA, early-adolescent WD diet exposure; WD-LA, late-adolescent WD diet exposure; NOIC, Novel Object in Context; PN, postnatal day.

#### 3.4. Early-but not late-adolescent WD consumption impairs HPC-dependent memory function in male rats after longer-term healthy diet intervention, whereas deficits in female rats were reversed by longer-term healthy diet intervention

In male rats, the initial memory deficits (Fig. 4A; P=0.0002 overall) persisted despite the longer healthy diet intervention in WD-EA rats only, whereas in WD-LA the deficits were reversed (Fig. 4B; P=0.0007 overall). Post hoc Tukey’s comparisons revealed that group WD-EA differed from both the CTL and WD-LA groups, and that the CTL and WD-LA groups did not differ from each other (CTL vs. WD-EA, P=0.0007; CTL vs. WD-LA, P=0.4541; WD-EA vs. WD-LA, P=0.0146). In females, however, the initial memory impairments (Fig. 4C; P=0.0017 overall) were reversed following the longer healthy diet intervention period during adulthood in both the WD-EA and WD-LA groups (Fig. 4D; P=0.1732). These NOIC results were consistent when examined as discrimination indices on day 3 and day 5 of the NOIC task (Supplementary Fig. S2; i.e., no differences in baseline discrimination index for the future novel object but differences in discrimination index on the test day for the time point after the longer healthy diet intervention period), and occurred in absence of any differences in total object exploration time on day 3 and day 5 of the NOIC task (Supplementary Fig. S2).

#### 3.5. Neither early-nor late-adolescent WD diet consumption alters object-novelty exploration, anxiety-like behavior, or locomotor activity

Because we previously did not find differences in object-novelty exploration (NOR), anxiety-like behavior (Zero Maze and Open Field), or locomotor activity (Open Field [distance travelled]) after WD diet consumption during the entire adolescent period (PN 26-56) in male rats (Hayes et al., 2023), we only examined these outcomes in females for the present work.

For NOR, there were no differences in object exploration index between the WD-EA, WD-LA, and CTL groups after either the shorter or longer healthy diet intervention periods (Fig. 5A-B; P=0.2605 and P=0.6280, respectively).

**Figure 5.**
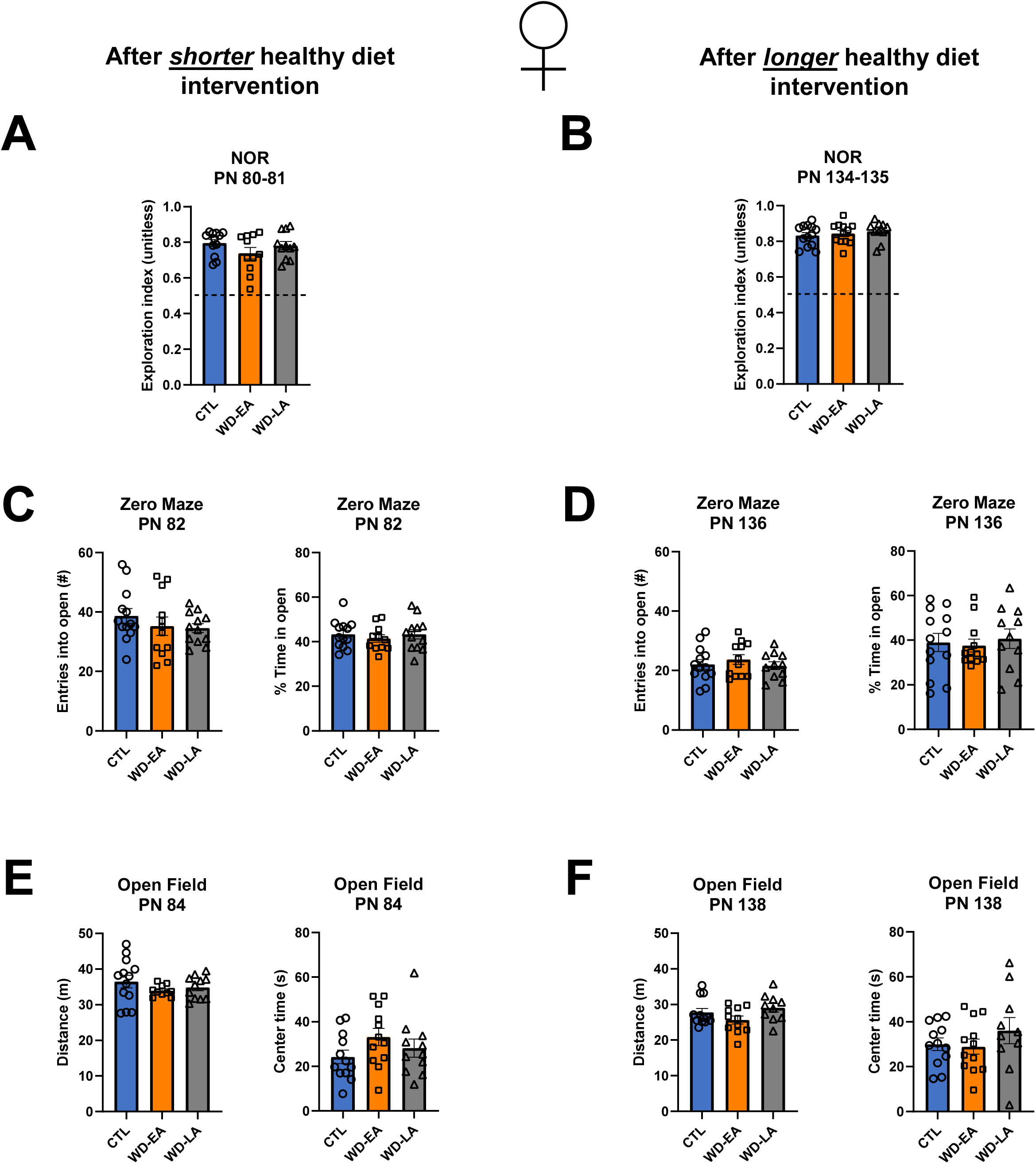
Early-adolescent (WD-EA) and late-adolescent (WD-LA) consumption of WD diet does not significantly affect hippocampus-independent novel object recognition memory function or anxiety-like behavior in female rats. (A) Perirhinal cortex-dependent object-novelty recognition assessed via NOR in female rats after early-adolescent (WD-EA) and late-adolescent (WD-LA) WD diet exposure and after the initial shorter healthy diet intervention period (one-way ANOVA with diet as the factor; P=0.2605). (B) Perirhinal cortex-dependent object-novelty recognition in female rats after early-adolescent (WD-EA) and late-adolescent (WD-LA) WD diet exposure and after the longer healthy diet intervention period (one-way ANOVA with diet as the factor; P=0.6280). (C) Anxiety-like behavior assessed via Zero Maze in female rats after early-adolescent (WD-EA) and late-adolescent (WD-LA) WD diet exposure and after the initial shorter healthy diet intervention period (entries into open: one-way ANOVA with diet as the factor; P=0.4446; percent time in open: one-way ANOVA with diet as the factor; P=0.7203). (D) Anxiety-like behavior assessed via Zero Maze in female rats after early-adolescent (WD-EA) and late-adolescent (WD-LA) WD diet exposure and after the longer healthy diet intervention period (entries into open: one-way ANOVA with diet as the factor; P=0.6381; percent time in open: one-way ANOVA with diet as the factor; P=0.8516). (E) Anxiety-like behavior and locomotor activity assessed via Open Field in female rats after early-adolescent (WD-EA) and late-adolescent (WD-LA) WD diet exposure and after the initial shorter healthy diet intervention period (distance: one-way ANOVA with diet as the factor; P=0.3860; center time: one-way ANOVA with diet as the factor; P=0.2335). (F) Anxiety-like behavior and locomotor activity assessed via Open Field in female rats after early-adolescent (WD-EA) and late-adolescent (WD-LA) WD diet exposure and after the longer healthy diet intervention period (distance: one-way ANOVA with diet as the factor; P=0.1213; center time: one-way ANOVA with diet as the factor; P=0.4298). Error bars represent ± standard error of the mean; n=10-13/group. ANOVA, analysis of variance; WD, cafeteria diet; WD-EA, early-adolescent WD diet exposure; WD-LA, late-adolescent WD diet exposure; NOR, Novel Object Recognition; PN, postnatal day.

For Zero Maze, we quantified the number of entries into the open arms as well as the percentage of time that rats spent in the open arms. Results revealed no differences in either of these outcomes after either the shorter or longer healthy diet intervention periods (Fig. 5C-D; P=0.4446 for entries into the open arms and P=0.7203 for percent time in the open arms after the shorter healthy diet intervention, P=0.6381 for entries into the open arms and P=0.8516 for percent time in the open arms after the longer healthy diet intervention).

Through the Open Field task, we assessed locomotor activity by quantifying the distance the rats travelled and gained additional insight into anxiety-like behavior by measuring the amount of time the rats spent in the brighter center zone of the box. There were no differences in either of these outcomes, regardless of experiencing a shorter vs. longer healthy diet intervention (Fig. 5E-F; P=0.3860 for distance and P=0.2335 for center time after the shorter healthy diet intervention, P=0.1213 for distance and P=0.4298 for center time after the longer healthy diet intervention).

### 4. Discussion

Given the widespread consumption of a Western diet (WD) and its link with cognitive dysfunction (Francis & Stevenson, 2013; Kanoski & Davidson, 2011; Kanoski et al., 2010), it is imperative to understand the critical periods of development during which WD consumption can induce long-lasting cognitive impairments that remain in adulthood. The present findings reveal that consumption of a WD in either early-or late-adolescence resulted in hippocampus (HPC)-dependent memory impairments after a shorter healthy diet intervention period (2-4 weeks). The extent to which these impairments were long-lasting after a prolonged healthy diet intervention during adulthood (10-12 weeks), however, was both sex- and adolescent window-specific. In male rats, consumption of WD in early-adolescence imparted long-lasting memory impairments that persisted despite a longer healthy diet intervention in adulthood, whereas initial memory deficits due to WD intake during late-adolescence were reversible following the longer dietary intervention. In contrast, in female rats, a longer healthy diet intervention reversed initial memory impairments due to WD consumption during either early-or late-adolescence. WD-induced memory impairments were not secondary to effects on body weight, anxiety-like behavior, or object-novelty recognition, as no group differences were observed in these outcomes for either sex.

Previous studies have investigated the effects of early life consumption of various versions of a WD on neurocognitive outcomes (Ferreira et al., 2018; Kaczmarczyk et al., 2013; Khazen et al., 2019; Lalanza et al., 2014), yet to our knowledge the present work is the first to reveal sex-specific and enduring effects of WD consumption tied to a specific window of adolescence. Kaczmarczyk et al. (2013) found that consumption of a high-fat diet (HFD) containing 60% kcal from fat (vs. a low-fat diet containing 10% kcal from fat) for 1- or 3-weeks beginning at PN 21 both impaired memory function in male C57BL/6J mice, and 1-week (but not 3-week) consumption increased anxiety-like behavior. However, in the same study, switching back to the low-fat diet for 1 week reversed the memory deficits in the 1-week group, whereas reversibility was not assessed in the 3-week group (Kaczmarczyk et al., 2013). These results indicate that a 1-week window from PN 21-28 is insufficient to impart long-lasting memory impartments, even in males. These findings align with the reversible impacts of HFD (60% kcal from fat) exposure for 1 week (PN 35-42) on hippocampal DCX^+^ structure and BDNF expression in rats found by Chiazza et al. (2021). Khazen and colleagues (2019) examined whether 1-week juvenile (PN 21-28) vs. adult (PN 60-67) consumption of a HFD (60% kcal from fat) vs. control diet (4% kcal from fat) affected HPC-dependent memory function in male Sprague Dawley rats. Their results revealed that this duration of exposure was sufficient to impair memory function in juvenile rats and actually improved memory function in adult rats (Khazen et al., 2019), but outcomes were not examined for reversibility later in life. Present results together with these previous findings collectively suggest that, in male rodents, 1-week WD exposure during early-adolescence is sufficient to impart HPC dysfunction, but that a ∼2-week exposure period may be needed for deficits to endure well into adulthood. Given the 1-3 week time-course of cell maturation in the HPC (Curlik et al., 2014; Kozareva et al., 2019), and that a progressive decrease in the rate of hippocampal neurogenesis occurs between adolescence and adulthood (He & Crews, 2007), the present results lend support for the hypothesis that effects of diet exposure during early-vs. late-adolescent windows on HPC memory function may be tied to neurogenesis. Additional research is needed to evaluate a potential connection between hippocampal neurogenesis and early-adolescent WD-induced memory impairments.

In addition to identifying early-adolescence as a critical period for long-lasting dietary influences on cognition in male rats, the present results highlight the need to consider sex as a biological variable in studying HPC vulnerabilities to dietary insults. The lack of long-lasting HPC-dependent memory impairments following 10-12 weeks of healthy diet intervention after early life WD exposure either during early-(PN 26-41) or late-adolescence (PN 41-56) in female rats aligns with previous research indicating a connection between HPC estradiol signaling and memory function in female rats (Chen et al., 2021; Yagi et al., 2023). Indeed, estrogen has previously been identified as a protective factor against aging (Norbury et al., 2003; Wise et al., 2001; Yang et al., 2020). Moreover, evidence suggests estrogen is also protective against WD-induced memory impairments in both rats and macaques (Estrada-Cruz et al., 2023; Scudiero & Verderame, 2017; Zimmerman et al., 2020). In addition to sex hormones, it is also possible that the gut microbiome is related to the sex-dependent results observed in the present study. While the microbiome was not examined in the present experiments, our previous work revealed that 30-day WD consumption during the entire adolescent period led to gut microbiota dysbiosis persisting after an adult healthy diet intervention in females, but not in males (Hayes et al., 2023; Tsan, Sun, et al., 2022). Future research is required to better understand whether the sex-dependent deviations in behavioral susceptibility to long-lasting WD-induced HPC dysfunction are functionally connected to changes in gut bacterial populations.

Although animals receiving WD diet in the present study generally consumed more energy (kcal) than their counterparts receiving standard chow at the same time, such increased intake did not lead to differences in body weight between groups, nor did they persistent when WD groups were returned to healthy standard chow. This could be related to the relatively short duration of time that the groups were exposed to WD diet, the fact that the adolescent rats were undergoing a period of rapid growth during which excess calories may be more readily used (Holliday, 1986), and/or due to altered energy expenditure more broadly (Jackman et al., 2010). While a hypercaloric diet has been linked with cognitive dysfunction (Kanoski et al., 2010; Morin et al., 2017; Reichelt et al., 2018), considering that both early- and late-adolescent WD consumption increased energy intake but both did not lead to long-lasting memory impairments, it is unlikely that caloric overconsumption is specifically tied to persistent HPC dysfunction. In our previous study in which female rats were given a WD diet during the entire adolescent period, there were no differences in energy intake between the WD group and controls, and the WD group nevertheless exhibited long-lasting memory impairments (Tsan, Sun, et al., 2022). Examining our present WD intake results in finer detail, there were no statistically significant differences in percentages of kcal consumed from the WD food components or percentages of macronutrients consumed between male WD-EA and WD-LA groups indicating that, in the context of a WD, long-lasting memory deficits may not be tied to greater consumption of certain high-fat high-sugar foods and/or macronutrients. This is further supported given that, in female rats, significant differences in percentage of kcal consumed from select WD components and macronutrients were observed, but in females the WD-induced memory impairments were reversible with longer term healthy diet intervention. Our previous and current findings together suggest that early life WD-associated HPC dysfunction is not based on elevated caloric consumption or greater consumption of a specific WD component or macronutrient, but future research is needed to verify this as well as to parse apart the potential role of altered energy expenditure.

Collectively, the present results identify early-adolescence as a critical window during which the HPC in male, but not female, rats is particularly vulnerable to WD-induced long-lasting HPC-dependent memory impairments that persist despite a 10-12 week healthy diet intervention. These findings offer valuable insights into the crucial influences of diet consumption in early life on cognition, with translational relevance for targeting dietary strategies to help prevent WD-induced cognitive dysfunction in a sex-specific manner.

## 5. Acknowledgments

The authors acknowledge the contributions of all the Kanoski Lab undergraduate researchers for their assistance with experimentation. Figure 1 was created with assistance from BioRender.com.

## 6. Author Contributions

The authors’ contributions were as follows: AMRH, LT, and SEK designed the research; AMRH, AEK, AA, KSS, MEK, JJR, ACN, and LT conducted the experiments; AMRH analyzed the data; AMRH and SEK wrote the manuscript with input from all authors; all authors read and approved the final manuscript.

## 7. Funding

This work was supported by the National Institute of Diabetes and Digestive and Kidney Diseases under grant DK123423 (awarded to SEK); the National Institute on Aging through a Postdoctoral Ruth L. Kirschstein National Research Service Award under grant F32AG077932 (awarded to AMRH); National Science Foundation Graduate Research Fellowships (separate awards to KSS and LT); and the National Institute of Diabetes and Digestive and Kidney Diseases through a Predoctoral Ruth L. Kirschstein National Research Service Award under grant DK13765655 (awarded to JJR).

## 8. Declarations of interest

The authors report there are no competing interests to declare.

## 9. Data availability statement

All data associated with this article are included in Supplemental Dataset 1. The authors confirm that the data supporting the findings of this study are available within the article and its supplementary materials.

## 10. Supplementary material

Supplementary Figure S1, Supplementary Figure S2, Supplementary Table S1, and Supplemental Dataset 1 are included as supplementary material for this article.

## Abbreviations

ANOVA: analysis of variance
BDNF: brain-derived neurotrophic factor
CTL: control
DCX^+^: doublecortin^+^
HFCS: high-fructose corn syrup
HFD: high-fat diet
HFHSD: high-fat high-sugar diet
kcal: kilocalories
HPC: hippocampus
NOIC: Novel Object in Context
NOR: Novel Object Recognition
NMR: nuclear magnetic resonance
PN: postnatal day
SEM: standard error of the mean
WD: Western diet
WD-EA: early-adolescent WD exposure
WD-LA: late-adolescent WD exposure.

---

**Supplementary Figure S1.**
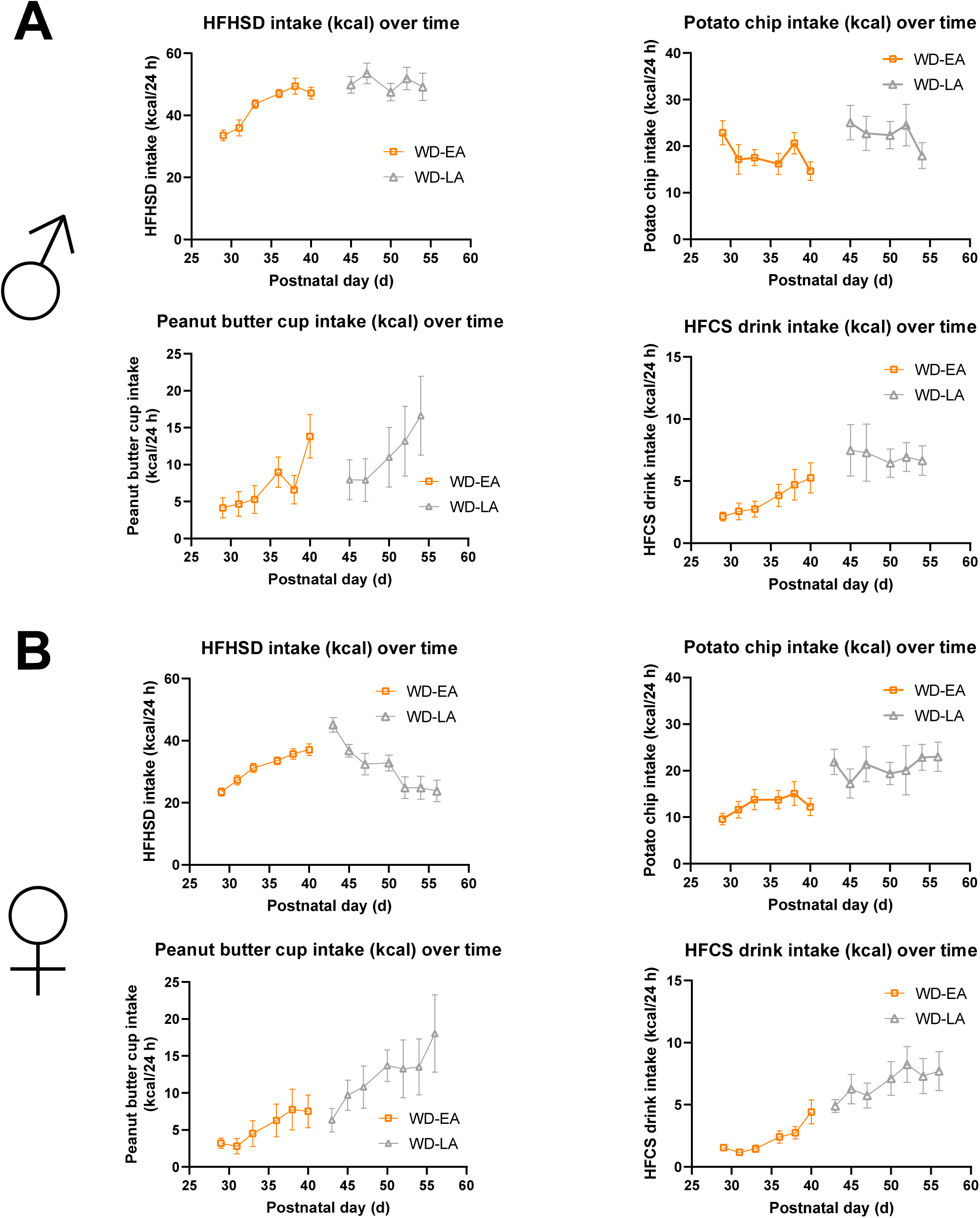
Energy consumed from the distinct diet components over time for early-adolescent (WD-EA) and late-adolescent (WD-LA) consumption of a WD diet. (A) Energy consumption over time for the high-fat high-sugar diet (HFHSD), potato chips, peanut butter cups, and high-fructose corn syrup (HFCS) in male rats (statistical analyses not performed). (B) Energy consumption over time for the high-fat high-sugar diet (HFHSD), potato chips, peanut butter cups, and high-fructose corn syrup (HFCS) in female rats (statistical analyses not performed). Error bars represent ± standard error of the mean; n=11-13/group. WD, cafeteria diet; WD-EA, early-adolescent WD diet exposure; WD-LA, late-adolescent WD diet exposure; HFCS, high-fructose corn syrup; HFHSD, high-fat high-sugar diet; kcal, kilocalories; PN, postnatal day.

**Supplementary Figure S2.**
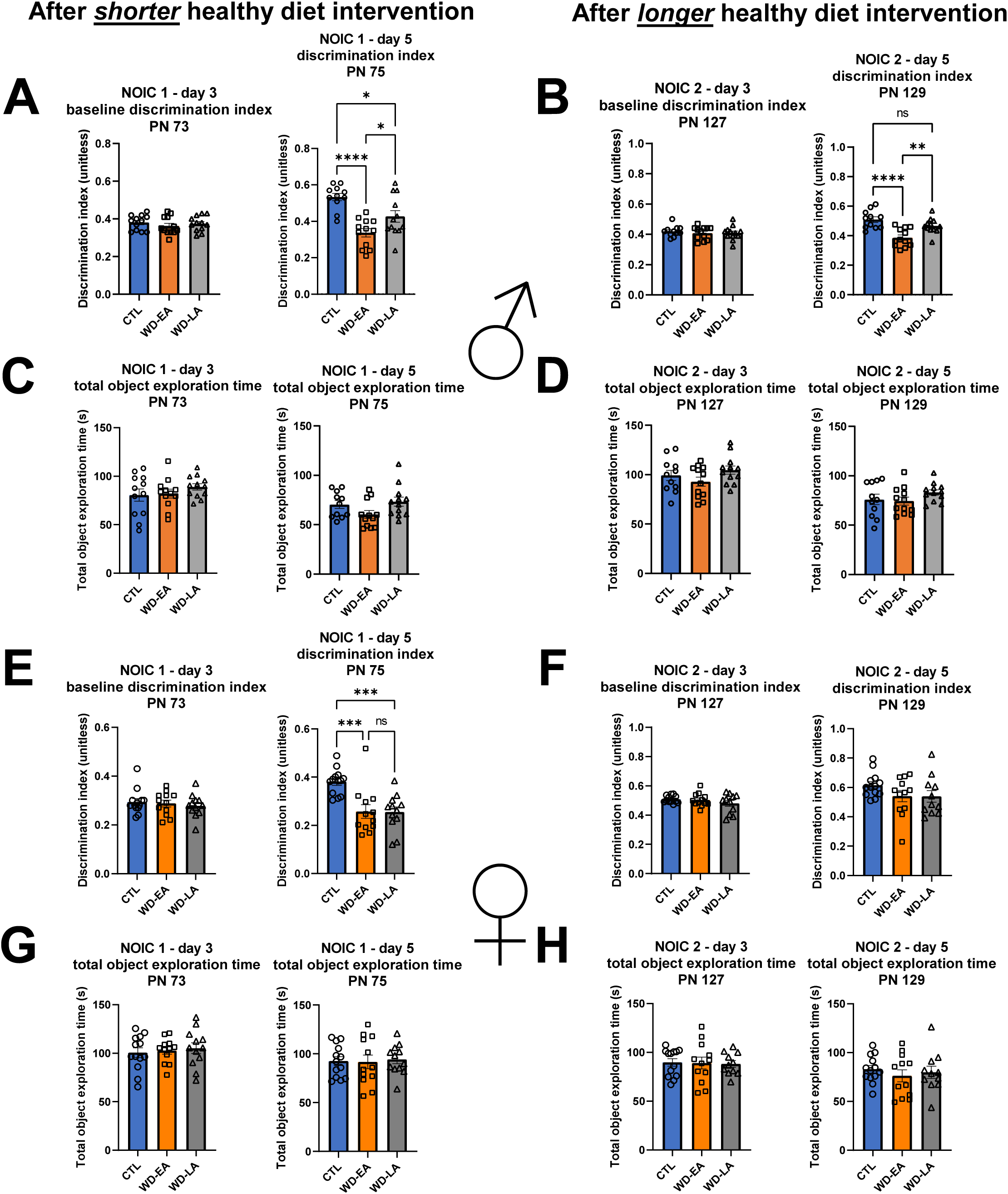
Early-adolescent (WD-EA) and late-adolescent (WD-LA) consumption of a cafeteria-style Western diet (WD) yielded differences in discrimination index of the novel object on the test day (day 5) but not the baseline discrimination index day (day 3), with no differences in total object exploration time on either day. (A) NOIC discrimination index for the future novel object (baseline, day 3) and novel object (test day, day 5) in male rats after early-adolescent (WD-EA) and late-adolescent (WD-LA) WD diet exposure and after the initial shorter healthy diet intervention period (baseline [day 3]: one-way ANOVA with diet as the factor, P=0.5973; test day [day 5]: one-way ANOVA with diet as the factor, P<0.0001 overall, post hoc Tukey’s multiple comparisons test: CTL vs. WD-EA, P<0.0001; CTL vs. WD-LA, P=0.0180; WD-EA vs. WD-LA, P=0.0487). (B) NOIC discrimination index for the future novel object (baseline, day 3) and novel object (test day, day 5) in male rats after early-adolescent (WD-EA) and late-adolescent (WD-LA) WD diet exposure and after the longer healthy diet intervention period (baseline [day 3]: one-way ANOVA with diet as the factor, P=0.7586; test day [day 5]: one-way ANOVA with diet as the factor, P<0.0001 overall, post hoc Tukey’s multiple comparisons test: CTL vs. WD-EA, P<0.0001; CTL vs. WD-LA, P=0.1996; WD-EA vs. WD-LA, P=0.0053). (C) NOIC total object exploration time in male rats after early-adolescent (WD-EA) and late-adolescent (WD-LA) WD diet exposure and after the initial shorter healthy diet intervention period (baseline [day 3]: one-way ANOVA with diet as the factor, P=0.4321; test day [day 5]: one-way ANOVA with diet as the factor, P=0.0858). (D) NOIC total object exploration time in male rats after early-adolescent (WD-EA) and late-adolescent (WD-LA) WD diet exposure and after the longer healthy diet intervention period (baseline [day 3]: one-way ANOVA with diet as the factor, P=0.2132; test day [day 5]: one-way ANOVA with diet as the factor, P=0.2752). (E) NOIC discrimination index for the future novel object (baseline, day 3) and novel object (test day, day 5) in female rats after early-adolescent (WD-EA) and late-adolescent (WD-LA) WD diet exposure and after the initial healthy diet intervention period (baseline [day 3]: one-way ANOVA with diet as the factor, P=0.7254; test day [day 5]: one-way ANOVA with diet as the factor, P=0.0002 overall, post hoc Tukey’s multiple comparisons test: CTL vs. WD-EA, P=0.0010; CTL vs. WD-LA, P=0.0008; WD-EA vs. WD-LA, P=0.9998). (F) NOIC discrimination index for the future novel object (baseline, day 3) and novel object (test day, day 5) in female rats after early-adolescent (WD-EA) and late-adolescent (WD-LA) WD diet exposure and after the longer healthy diet intervention period (baseline [day 3]: one-way ANOVA with diet as the factor, P=0.3399; test day [day 5]: one-way ANOVA with diet as the factor, P=0.1974). (G) NOIC total object exploration time in female rats after early-adolescent (WD-EA) and late-adolescent (WD-LA) WD diet exposure and after the initial shorter healthy diet intervention period (baseline [day 3]: one-way ANOVA with diet as the factor, P=0.7882; test day [day 5]: one-way ANOVA with diet as the factor, P=0.9456). (H) NOIC total object exploration time in female rats after early-adolescent (WD-EA) and late-adolescent (WD-LA) WD diet exposure and after the longer healthy diet intervention period (baseline [day 3]: one-way ANOVA with diet as the factor, P=0.9724; test day [day 5]: one-way ANOVA with diet as the factor, P=0.6685). Error bars represent ± standard error of the mean; n=11-13/group; *P<0.05, **P<0.01, ***P<0.001, ****P<0.0001. ANOVA, analysis of variance; WD, cafeteria diet; WD-EA, early-adolescent WD diet exposure; WD-LA, late-adolescent WD diet exposure; NOIC, Novel Object in Context; PN, postnatal day.

## Notes

### Competing Interest Statement

The authors have declared no competing interest.

